# Are the adverse effects of stressors on amphibians mediated by their effects on stress hormones?

**DOI:** 10.1101/165282

**Authors:** Caitlin R. Gabor, Sarah A. Knutie, Elizabeth A. Roznik, Jason R. Rohr

## Abstract

Adverse effects of anthropogenic changes on biodiversity might be mediated by their impacts on the stress response of organisms. To test this hypothesis we crossed exposure to metyrapone, a synthesis inhibitor of the stress hormone corticosterone, with exposure to the herbicide atrazine and the fungal pathogen *Batrachochytrium dendrobatidis (Bd)* to assess whether the effects of these stressors on tadpoles and post-metamorphic frogs were mediated by corticosterone. Metyrapone countered atrazine-and Bd-induced corticosterone elevations. However, atrazine-and *Bd*-induced reductions in body size were not mediated by corticosterone because they persisted despite metyrapone exposure. Atrazine lowered *Bd* abundance without metyrapone but increased *Bd* abundance with metyrapone for tadpoles and frogs. In contrast, atrazine reduced tolerance of *Bd* infections because frogs exposed to atrazine as tadpoles had reduced growth with *Bd* compared to solvent controls; this effect was not countered by metyrapone. Our results suggest that the adverse effects of atrazine and *Bd* on amphibian growth, development, and tolerance of infection are not mediated primarily by corticosterone. Instead, these effects are likely a function of energy lost from atrazine detoxification, defense against *Bd*, or repair from damage caused by atrazine and *Bd*. Additional studies are needed to evaluate how often the effects of anthropogenic stressors are mediated by stress hormones.

## Introduction

We are now in the age of the Anthropocene, a time when human activity is the dominant influence on the environment and biodiversity (Dirzo et al. 2014). Many anthropogenic factors, such as chemical contaminants and introduced pathogens, can function as stressors by elevating or dysregulating glucocorticoid “stress hormones” in vertebrates (Gabor et al. 2015; Larson et al. 1998; McMahon et al. 2017). In turn, these interactions can have profound and enduring effects on the health of organisms, especially when exposure occurs during early-life stages (Boekelheide et al. 2012; Martin et al. 2010; Rohr et al. 2013). For example, early-life exposure to stress hormones during key developmental stages can permanently alter the functionality of the hypothalamic-pituitary-adrenal (HPA; HPI-interrenal in amphibians) axis, which in turn can alter the immune system into adulthood (Martin et al. 2010; Matthews 2002; Rohr et al. 2013). Thus, even though many physiological responses to stress can be adaptive (Boonstra 2013), many of the adverse effects of the Anthropocene on biodiversity might be mediated by the stress physiology of organisms. However, this hypothesis has not been thoroughly tested because no studies to date have crossed anthropogenic stressors with compounds that inhibit the synthesis of stress hormones. Such a study would help determine whether stress hormones are largely responsible for adverse effects of anthropogenic stress.

Amphibians are the most threatened class of vertebrates on the planet (Stuart et al. 2004) and represent a taxon extensively impacted by activities dominating the Anthropocene. Anthropogenic factors have been implicated in amphibian declines, including environmental pollutants, infectious diseases, and their interactions (Hayes et al. 2010; Jones et al. 2017; Rohr and McCoy 2010). For instance, chytridiomycosis, the disease caused by the fungal pathogen, *Batrachochytrium dendrobatidis (Bd)*, has caused major declines and possibly extinctions of hundreds of amphibian species in the last half century (Wake and Vredenburg 2008).

Amphibians employ two defense strategies against *Bd* that can be impacted by anthropogenic factors and associated stress hormones. Resistance strategies, such as innate and adaptive immune responses (McMahon et al. 2014; Rollins-Smith et al. 2011), prevent or clear *Bd* infections, whereas tolerance strategies minimize the fitness consequences of infection, such as mechanisms that enhance repair from parasite damage (Råberg et al. 2009; Rohr et al. 2010).

The immune system in vertebrates is partly modulated by glucocorticoids, such as corticosterone, the main glucocorticoid related to stress in amphibians. Corticosterone and other stress hormones, such as cortisol, are regularly used to assess overall stress and health of wild animal populations and to direct wildlife management (reviewed by Busch and Hayward 2009; Sheriff et al. 2011). For example, chronically-elevated corticosterone can accelerate metamorphosis and decrease amphibian growth, development, and immunity, the latter of which can increase infectious disease risk (Denver 2009; Rollins-Smith 1998; Warne et al. 2011). Chemical contaminants, such as the herbicide atrazine, the second most commonly used pesticide in the US (Kiely et al. 2004), and pathogens, such as *Bd*, can elevate corticosterone (Gabor et al. 2015; McMahon et al. 2017; Peterson et al. 2013; Searle et al. 2014). Contaminants have similar negative effects on amphibians as chronic corticosterone, such as reduced immunity and growth (Larson et al. 1998; McMahon et al. 2013a; Rohr et al. 2013). This suggests that the negative effects of contaminants might be mediated by corticosterone. For example, early-life exposure to atrazine reduces amphibian growth and development and, despite not affecting amphibian resistance to *Bd*, it reduced tolerance of *Bd* infections, thus increasing Bd-induced mortality (Rohr et al. 2013).

Using a series of experiments where we inhibited corticosterone synthesis in Cuban treefrog *(Osteopilus septentrionalis)* using the compound metyrapone, we explore whether the effects of atrazine and *Bd* on amphibian growth and development, and the effects of atrazine on amphibian resistance and tolerance of *Bd* were mediated by their effects on amphibian corticosterone. We hypothesized that any effects of atrazine and *Bd* exposure on growth, development, and survival would be at least partly mediated by their effects on corticosterone. We also hypothesized that atrazine would affect amphibian-Bd interactions by altering amphibian resistance and/or tolerance of *Bd* and that this too would be at least partly mediated by corticosterone. Importantly, if corticosterone mediates any of these effects, then corticosterone levels should be correlated with these responses and metyrapone should counteract some or all of the effects of atrazine and *Bd*.

## MATERIALS AND METHODS

We collected multiple clutches of tadpoles of *O. septentrionalis* in August 2014 from the Botanical Gardens of the University of South Florida (N 28°03.537’ W 082°25.410’). We maintained them in the lab for at least a week until the majority reached Gosner developmental stage 35 (Gosner 1960). All tadpoles were fed a mixture of fish food and spirulina suspended in agar *ad libitum* and were maintained at 21°C with a 12 h light cycle. We noted tadpole survival daily.

### Baseline corticosterone and stress responses

To determine the physiological range of corticosterone in *O. septentrionalis* tadpoles, we quantified the release rate of water-borne corticosterone from undisturbed (“baseline”) tadpoles (n = 18) and we obtained the stress response from naturally stressed tadpoles (n = 15) through gentle “agitation”. Given that these methods are already well documented, these methods are presented in the Supplemental Material.

### Experimental overview

Our experiment consisted of three stages (Fig. 1). First, tadpoles were exposed to fully crossed atrazine and metyrapone (corticosterone synthesis inhibitor) treatments for six days. Metyrapone has been used in other studies with amphibians to explore the effects of corticosterone on stress responses (Glennemeier and Denver 2002a; Glennemeier and Denver 2002b), specifically the stress response to predation (Middlemis Maher et al. 2013; Neuman-Lee et al. 2015), and has not been found to change amphibian behavior (Hossie et al. 2010). Second, after placing tadpoles in fresh untreated water, we challenged half the amphibians with *Bd* or not. Third, we challenged the other half of tadpoles, once they were post-metamorphic frogs, with *Bd* or not. We obtained water-borne corticosterone release rates from tadpoles after the six days of atrazine exposure and again after a week of *Bd* exposure.

**Fig. 1.**
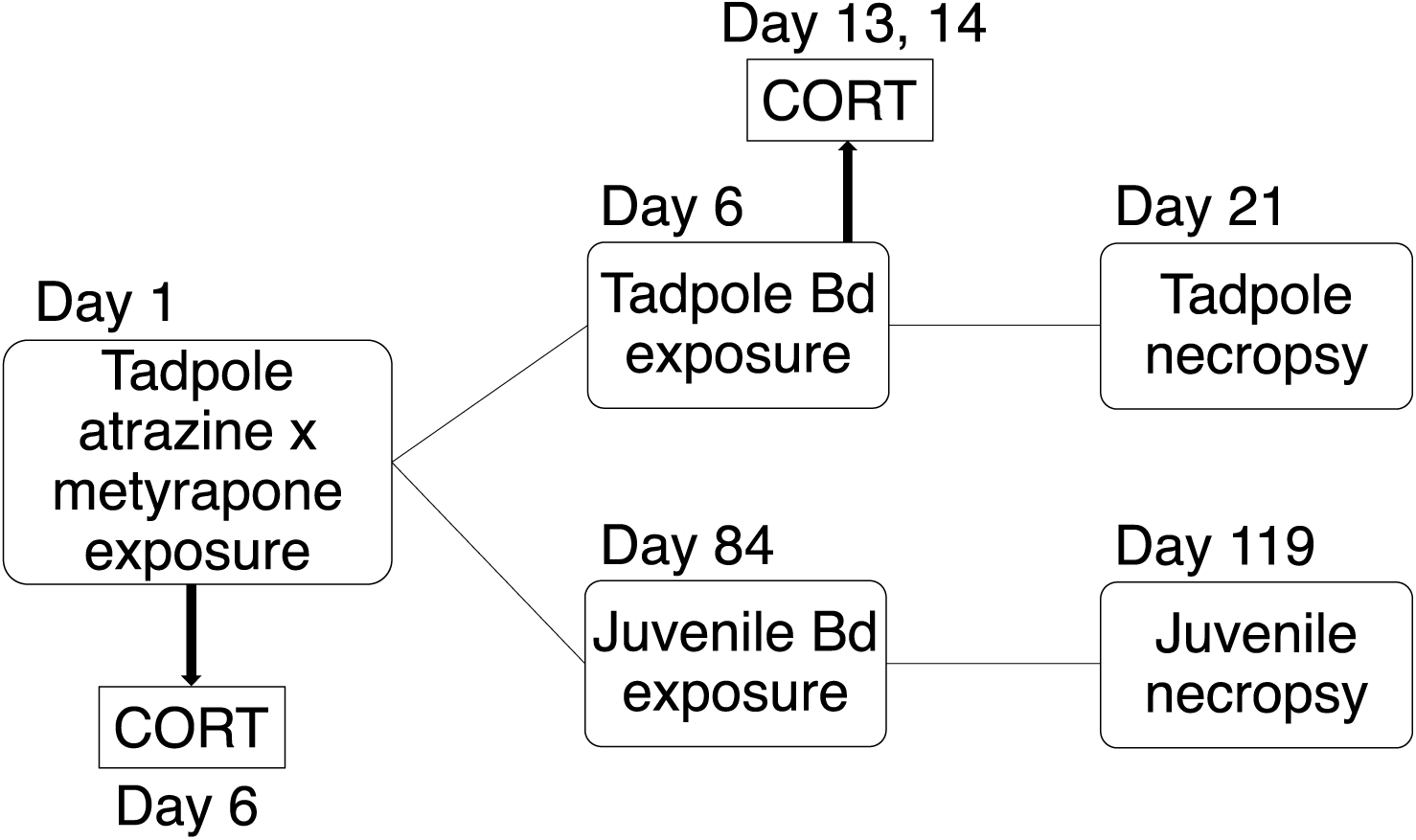
Flow chart showing the experimental design for exposing tadpoles of *Osteopilus septentrionalis* (n = 10 replicates, 16 tadpoles per replicate per treatment) to atrazine and metyrapone (corticosterone synthesis inhibitor) followed by movement to new tanks and exposure to *Batrachochytrium dendrobatidis (Bd)* or not. Half of the tadpoles from the first chemical exposure were reared to post-metamorphic (juvenile) frogs and then exposed to *Bd* (or not). All were euthanized for further analysis.

### Atrazine and metyrapone exposure in tadpoles

On the first day of our experiment, we filled forty 12-l tanks with 8 l of water from a pond at Trout Creek Park, FL (N28°092250’, W082 °348083’) that was free of tadpoles and was not exposed to agricultural runoff (i.e., no measurable level of atrazine, see atrazine measurements below). We assigned 16 *O. septentrionalis* tadpoles haphazardly to each tank. We randomly assigned each tank to one of four exposure treatments: (1) the estimated environmental concentration (EEC) of atrazine (200 μg l^-1^; Chemservice, West Chester, PA; technical grade, purity more than 98%) dissolved in 120 μl of ethanol (n = 10), (2) 110 μM of metyrapone (Sigma Chemical Co. # M2696; St. Louis, MO) dissolved in 120 μl of ethanol, (n = 10), (3) the EEC of atrazine and 110 μM of metyrapone jointly dissolved in 120 μl of ethanol (n = 11), and (4) only 120 μl of ethanol (n = 10) as a control. We used 110 μM of metyrapone because this level reduced whole body corticosterone in tadpoles by >50% (Glennemeier and Denver 2002b) and, over the short term (weeks), exposure is believed to be non-toxic (Glennemeier and Denver 2002c). Previous work did not detect effects of ethanol on any measured trait, and thus a water control was not included (reviewed by Rohr et al. 2013). Tadpoles were exposed to these treatments for six days.

The targeted nominal concentration of atrazine was 200 ugl^-1^, the EEC based on US Environmental Protection Agency GENEEC v2 software. The EEC is the concentration estimated to enter a standardized farm pond at a standardized distance from an application site given the chemical properties of the pesticide; thus, it is an ecologically relevant concentration. To quantify actual atrazine concentrations, water samples were taken from each of the 40 tanks 1 h after dosing and atrazine was quantified using the Abraxis ELISA microtiter plate kit (Abraxis LLC, Warminster, PA). Mean (± 1 SE) atrazine concentration was 178.2 ± 7.8 μg l^-1^. All atrazine values for the non-atrazine exposed tanks were below the detection limit of 0.06 μg l^-1^ (this is the level in the pond water). We re-dosed each tank with 110 μM of metyrapone every third day (following Hossie et al. 2010). We did not re-dose with atrazine because its half-life is on the order of weeks and Rohr et al. (2004) found no detectable breakdown of atrazine over seven days under similar conditions.

After tadpoles were exposed to their respective treatment, we obtained water-borne corticosterone (6d corticosterone) from two tadpoles per replicate (80 total) by placing tadpoles individually in 250ml beakers filled with 75ml of clean water for 1 hour (following Gabor et al. 2016). We then removed them and measured their mass and snout-vent-length (SVL). Gosner stage was not measured because the same tadpoles were used throughout the experiment but we noted that none had obvious limb buds. Water samples were frozen at-20°C immediately after collection until ready to be thawed for extraction; this method does not affect corticosterone levels (Ellis et al. 2004).

### *Bd* exposure in tadpoles

Immediately following the collection of corticosterone on day 6, eight tadpoles were removed from their original tank and were evenly divided between two 6-l plastic shoeboxes with 2 l of fresh pond water (for a total of 80 containers) for the *Bd* exposure stage of the experiment. *Bd* isolate SRS812 was cultured following the methods of McMahon et al. (2013a). Half of these shoeboxes received a 6-ml inoculum containing 7×104 *Bd* zoospores ml^-1^ in deionized (DI) water and the other half received an inoculum that was identical to the *Bd* inoculum but was free of *Bd* (i.e., we washed clean agar plates with DI water). We re-exposed all tadpoles to *Bd* (2 ml of 3×105 zoospores ml^-1^) or DI water three days later and maintained the tadpoles in these boxes until day 21.

On day 13 and 14 (seven and eight days after exposure to *Bd*), we collected water-borne corticosterone samples (13,14d corticosterone) from two tadpoles of each replicate to examine the effect of *Bd* exposure on individual corticosterone release rates (half the tadpoles each day using the same methods as described above). All other tadpoles were placed in the same size containers with the same amount of water to control for any effects of handling. All tadpoles were returned to their original tank after being placed in individual beakers. Twenty-one days after exposure to *Bd* or not, we swabbed each surviving tadpole for *Bd* by passing a sterile rayon swab along its mouthparts (eight strokes horizontally, and eight strokes vertically). We then euthanized tadpoles with an overdose of MS-222. We recorded their mass, SVL, and Gosner stage, and preserved them in 70% ethanol. We used quantitative PCR (described by Boyle et al. 2004) to quantify *Bd* abundance taken from up to two tadpoles per Bd-exposed tank (depending on survival, n = 69 total), and a total of 10 tadpoles (each from separate tanks) that were not exposed to *Bd.*

### *Bd* exposure in post-metamorphic frogs

Individuals that were not used in the *Bd* experiment as tadpoles remained in their original tanks until they metamorphosed. After the six-day chemical treatment, freshwater free of atrazine was provided every other week. Tadpole survival and day of metamorphosis (all four limbs had emerged; Gosner stage 42) were recorded daily. Upon metamorphosis, individuals were removed from the tanks and placed in cups (6 cm high×12 cm diameter) with moist *Sphagnum* sp. moss. The post-metamorphic frogs were maintained in the laboratory (12h light cycle, 22°C) and fed *ad libitum* vitamin-and mineral-dusted crickets twice per week. Frog survival was recorded daily. A week prior to *Bd* exposure, body mass was recorded for each individual. On day eighty-four of the experiment and approximately one month after most of the tadpoles metamorphosed, half the surviving post-metamorphic frogs from each tank were randomly assigned to receive *Bd* (isolate SRS812) and the other half received the control (each tank had 1-2 frogs exposed to each treatment depending on survival). Post-metamorphic frogs were exposed to *Bd* or a control solution (everything but *Bd)* by pipetting 1 ml of 6×104 zoospores ml^-1^ onto the frog’s dorsal side. Excess inoculum remained in each frog’s plastic container, which contained moist sterile *Sphagnum* moss. Survival was monitored daily for 5 weeks. Frogs were also weighed weekly and swabbed at two and three weeks after *Bd* exposure. *Bd* from the swabs was quantified using the methods described above. We then euthanized individuals with an overdose of MS-222 and preserved them in 70% ethanol for further processing.

### Hormone extraction and validation

We obtained all water-borne hormone samples between 0830-1500 h. We extracted water-borne hormones following (Gabor et al. 2016). We re-suspended the dried hormone residue in 260 μl enzyme-immunoassay (EIA) buffer (provided by Cayman Chemicals Inc., Ann Arbor, MI, USA) and we further diluted all samples to 1:2. We measured corticosterone in duplicate using a corticosterone EIA kit (Cayman Chemicals Inc.) on a spectrophotometer plate reader set to 405 nm (BioTek ELX800). Because tadpoles were placed in clean water, no chemicals from the exposure stage interacted with the corticosterone assay. See Supplemental material, Methods for validation of the water-borne corticosterone collection method from *O. septentrionalis* on EIA plates and intra-plate and plate-independent variation.

### Statistical analyses

Statistical analyses were conducted using R statistical software (R Core Team 2013) and all probability values were calculated using type II sums of squares. We used a general linear mixed-effects model (GLMM; nlme package) to test for all main and interactive effects of atrazine and metyrapone exposure on log-transformed 6d corticosterone release rates (adjusted for mass (pg g^-1^ h^-1^)) with tank as a random factor and EIA test plate and SVL as covariates. We also used the nlme package to test for all main and interactive effects of atrazine, metyrapone, and *Bd* treatments on 13,14d corticosterone release rates (pg g^-1^ h^-1^) using least trimmed squares with EIA test plate and SVL as covariates and tank as a random effect. We used a GLMM to test for the main and interactive effects of atrazine and metyrapone on the number of days to metamorphosis, treating tank as a random effect. To evaluate the effects of treatments on frog growth after metamorphosis, we conducted two-way MANOVAs (atrazine, metyrapone, atrazine×metyrapone) using log-transformed mass and SVL as response variables. We only present the mass data because mass and SVL were highly correlated. To determine the effect of treatments on tadpole and post-metamorphic frog survival, we conducted a mixed effects Cox proportional hazards survival analysis using the coxme function in the coxme R package.

To test for the effects of treatments on *Bd* resistance, we used the nlme package and lme function to test for the fully crossed effects of life stage (tadpole vs frog), atrazine, and metyrapone on log-transformed *Bd* abundance with sampling time (individuals sampled before and after metamorphosis) nested in tank as random effects. To test for effects on tolerance of *Bd*, we again used the lme function to test for the fully crossed effects of life stage, atrazine, metyrapone, and log *Bd* abundance on the percent mass change of each amphibian (standardized) with sampling time nested in tank as random effects. As a reminder, tolerance is measured as the slope of the relationship between pathogen load and a fitness proxy. Thus, a treatment with a more negative slope is less tolerant of infection (Råberg et al. 2007). When higher order interactions, covariates, or blocking factors were not significant, they were dropped from the statistical model. Finally, we performed Pearson tests to explore the correlations between 6d corticosterone and mass, development, time to metamorphosis, *Bd*, survival, resistance and tolerance for tadpoles and post-metamorphic frogs (all based on tank means). All parametric analyses met the underlying assumptions and we applied an alpha of 0.05.

## Results

### Baseline corticosterone and stress responses

We found that baseline corticosterone release rates were significantly lowe than the corticosterone release rates of agitated (stress response), and atrazine-exposed tadpoles. See Supplemental material, Results

### Effects on tadpoles and metamorphosis

Atrazine elevated corticosterone on experimental day 6 and *Bd* alone elevated corticosterone on experimental day 13 and 14, but metyrapone countered these increases in corticosterone induced by both factors (metyrapone×atrazine: χ^2^ = 8.20, d.f. = 1,*p* = 0.004; metyrapone×*Bd*: *χ*^2^ = 3.87, d.f. = 1, *p* = 0.049; Fig. 2a, b). Tadpoles exposed to atrazine alone had higher corticosterone than all other treatments *(p* < 0.05, Tukey's HSD). However, metyrapone plus atrazine did not have significantly different amounts of corticosterone than the control treatment (no atrazine or metyrapone) *(p* > 0.05, Tukey's HSD; Fig. 2a), indicating that the metyrapone successfully inhibited corticosterone synthesis. Tadpoles exposed to *Bd* alone had higher corticosterone than the control treatment (not exposed to *Bd)* and those exposed to *Bd* plus metyrapone *(p* < 0.05, Tukey's HSD), but metyrapone alone did not have significantly different amounts of corticosterone than these two treatments (Fig. 2b). Although atrazine elevated corticosterone during the atrazine exposure period (at day 6), this effect was not detectable after 15 days in atrazine-free water (Supplemental Table 1). There was no significant relationship between mean experimental corticosterone on day 6 and mean log *Bd* load of tadpoles and post-metamorphic frogs (Supplemental Table 1).

**Fig. 2.**
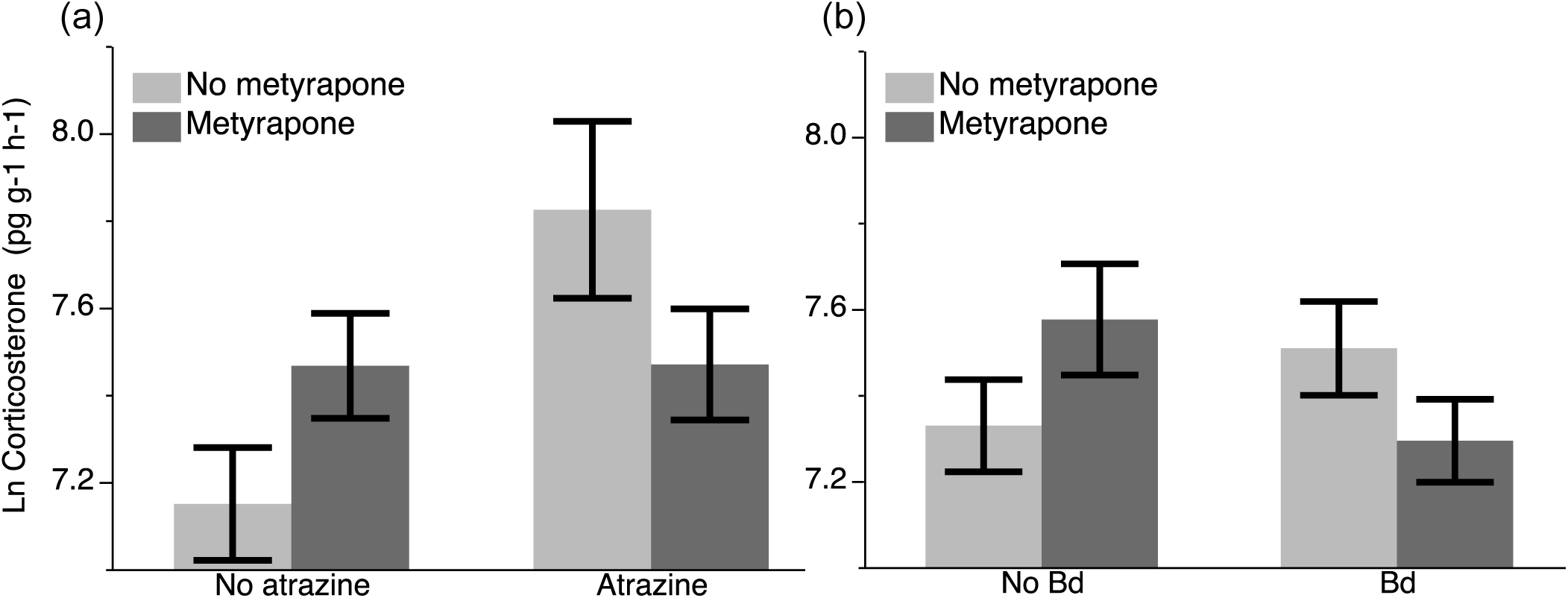
Ln corticosterone (± 1 SE) for tadpoles of *Osteopilus septentrionalis* after: (a) six days (6d) of exposure to metyrapone (CORT synthesis inhibitor) and atrazine or not, and (b) seven or eight days of exposure (13,14d) to metyrapone and *Batrachochytrium dendrobatidis (Bd)* or not. Both the atrazine×metyrapone and the metyrapone×*Bd* interactions are significant *(p* < 0.05). See Results for statistics.

Exposure to atrazine reduced tadpole body size one week into the experiment (MANOVA on SVL and mass: *F*_1,36_ = 3.60, *p* = 0.04; Fig. 3a), but metyrapone had no effect on body size at this time (metyrapone: *F*_1,36_ = 0.00, *p* = 1.00; atrazine×metyrapone: *F*_1,36_ = 1.32, *p* = 0.25; Fig. 3b). *Bd* exposure had no significant effects on mass or SVL *(p* > 0.05). Tadpole survival (% mortality) was not significantly affected by atrazine *(χ^2^* = 0.30, d.f. = 1, *p* = 0.86) or metyrapone (*χ*^2^ = 0.30, d.f. = 1, *p* = 0.58; Fig. 3c). Tadpole survival was weakly affected by an interaction between *Bd* and metyrapone; metyrapone tended to decrease days alive (survival) in the absence of *Bd* but increased it slightly in the presence of *Bd (χ^2^* = 3.87, d.f. = 1, *p* = 0.05; Fig. 4a). Time to metamorphosis was not significantly affected by atrazine, metyrapone, or their interaction (atrazine: *χ*^2^ = 0.00, d.f. = 1, *p* = 0.99, metyrapone: *χ*^2^ = 0.10, d.f. = 1, *p* = 0.75, atrazine×metyrapone: *χ*^2^ = 0.00, d.f. = 1, *p* = 0.95; Supplemental Fig. 1). Similarly, *Bd* had no effect on time to metamorphosis *(p* > 0.05). See Supporting Information, Results for effects of atrazine, metyrapone, and *Bd* on Gosner stage. Corticosterone on experimental day 6 was not correlated significantly with tadpole mass, time to metamorphosis, or survival *(p* > 0.05; see Supplemental Table 1).

**Fig. 3.**
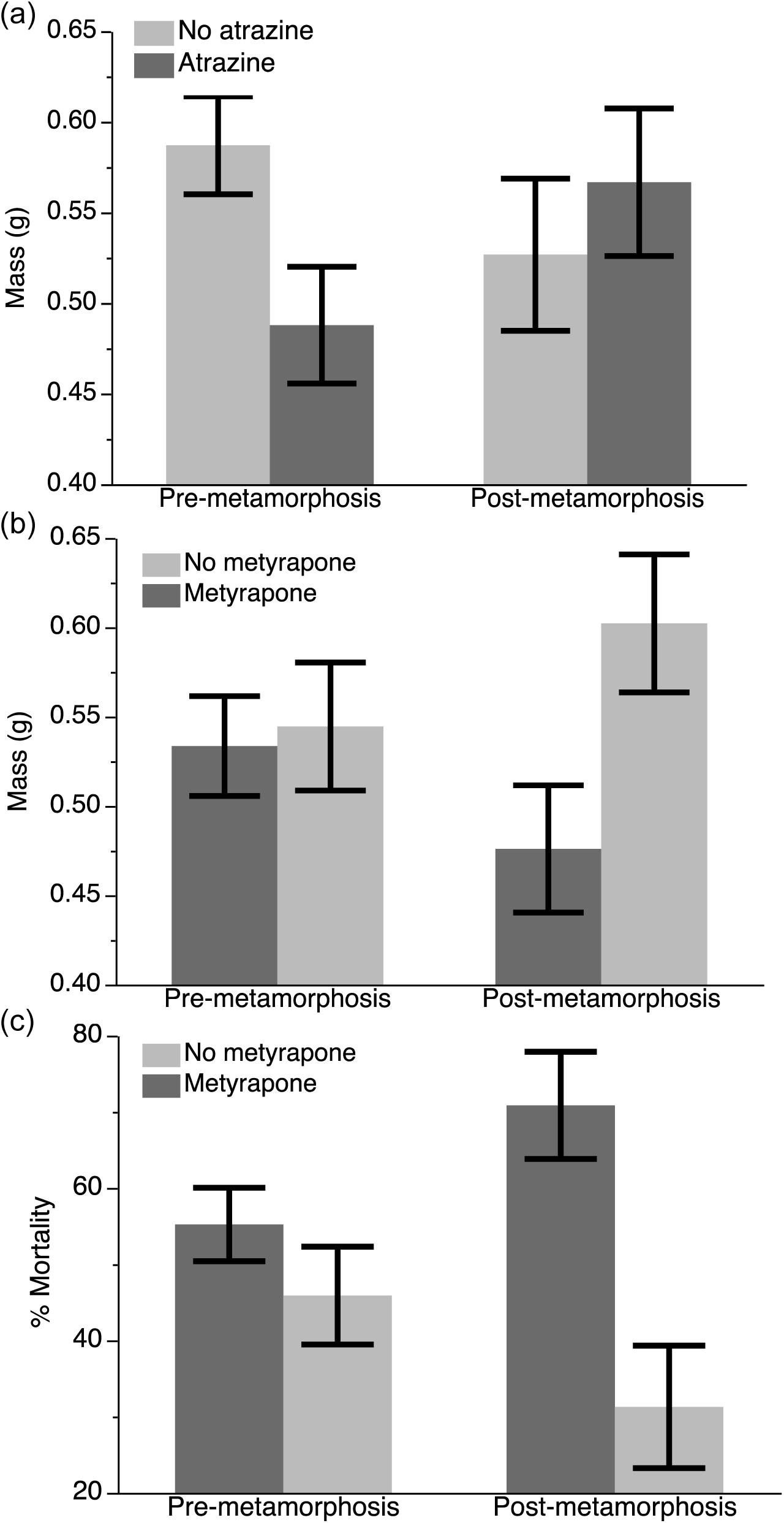
The mean mass (g) (± 1 SE) for pre-and post-metamorphic frogs with exposure to: (a) atrazine (n = 9 pre, 8 post) or (b) metyrapone (n = 11 pre, 5 post; corticosterone synthesis inhibitor). The mass change is significantly different for pre-metamorphic tadpoles exposed to atrazine or not and for post-metamorphic frogs exposed to metyrapone or not *(p* < 0.05). (c) Mean percent mortality (± 1 SE) of pre-(n = 41) and post-metamorphic (n = 41) frogs after exposure to metyrapone. The percent mortality is significantly different for post-metamorphic frogs exposed to metyrapone or not *(p* < 0.05). See Results for statistics.

**Fig. 4.**
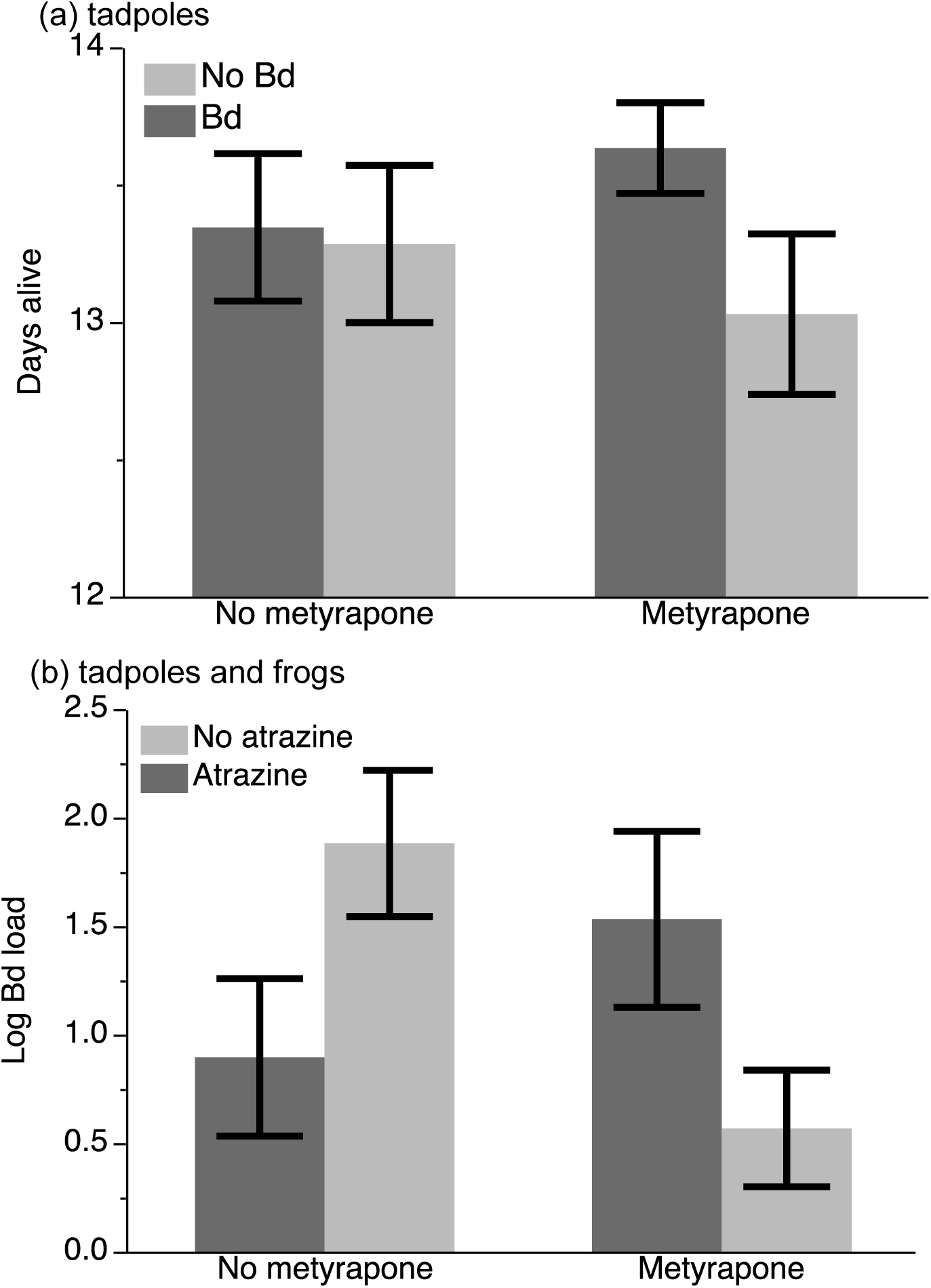
(a)The interactive effect of six days of metyrapone (corticosterone synthesis inhibitor) and subsequent *Batrachochytrium dendrobatidis (Bd)* exposure (15 days) on mean days alive (± SE) for tadpoles. The *Bd* main effect and metyrapone×*Bd* interaction are significant *(p* < 0.05). (b) The interactive effect of metyrapone (n = 22 no atrazine, n = 17 atrazine) and atrazine (n = 14, n = 16 no atrazine) on mean resistance (i 1 SE) to the fungal pathogen *Bd* (measured as log *Bd* abundance averaged across the tadpole and post-metamorphic frog exposure periods is significant *(p* < 0.05) for *Osteopilus septentrionalis.* Because there was no interaction with life stage we combined data across life stages. See Results for statistics.

### Effects on post-metamorphic frogs

For post-metamorphic frogs, prior exposure to atrazine did not have a significant effect on survival (χ^2^ = 0.02, d.f. = 1, *p* = 0.88) or body size (MANOVA, *F*_2,64_ = 1.82, *p* = 0.17) and there were no interactive effects of atrazine and metyrapone on these responses (*F*_2,64_ = 0.07, *p* = 0.93). Thus, atrazine reduced size before metamorphosis when measured soon after the chemical exposure but this size difference did not persist post-metamorphosis (stage×atrazine: *F*_1,61_ = 5.48, *p* = 0.02; Fig. 3a). In contrast, metyrapone reduced mass (*F*_2,64_ = 4.03, *p* = 0.02) and survival (χ^2^ = 15.37, d.f. = 1,*p* < 0.001) after metamorphosis, but did not significantly affect these responses before metamorphosis (Fig. 3b, c), indicating that there were delayed effects of metyrapone. Exposure to *Bd* did not significantly affect frog survival (days alive) after metamorphosis *(p* > 0.05 for all effects including *Bd).* Once again, corticosterone in tadpoles at experimental day 6 did not significantly correlate with post-metamorphic mass or survival *(p* > 0.05, see Supplemental Table 1).

### Effects of atrazine and metyrapone on resistance and tolerance of ***Bd* in both life stages**

Of the Bd-exposed tadpoles, 21 of 70 (30%) tested positive for *Bd* with loads ranging from 625 - 52,045 zoospores; we confirmed that control tadpoles were not infected with *Bd.* Of the post-metamorphic frogs exposed to *Bd*, 14 of 34 (41%) became infected, with loads ranging from 154-10,010 zoospores.

Atrazine and metyrapone significantly affected resistance to *Bd.* Atrazine lowered *Bd* abundance in the absence of metyrapone, but elevated *Bd* abundance in the presence of metyrapone (atrazine×metyrapone χ^2^ = 9.51, d.f. = 1, *p* = 0.002; Fig. 4b), and this effect was consistent across life stages (all effects including life stage had*p* > 0.22).

Most frogs survived the atrazine and *Bd* treatments and thus we focused on the ability of frogs to maintain or increase body size in the face of infection as our measure of tolerance. Both atrazine and metyrapone significantly affected tolerance of *Bd*, but this relationship was dependent on life stage of the individual frog. Metyrapone did not significantly affect tolerance of *Bd* as tadpoles, but reduced tolerance of post-metamorphic frogs. Frogs previously exposed to metyrapone lost more weight per *Bd* zoospore than frogs not previously exposed to metyrapone (stage×metyrapone×*Bd* load: /2 = 4.26, d.f. = 1, *p* = 0.04; Fig. 5a). This is consistent with the delayed adverse effects of metyrapone observed for growth and survival (Fig. 3b, c). Atrazine reduced tolerance of *Bd* both before and after metamorphosis. In the full statistical model, there was no evidence of any interactions between atrazine and life stage or atrazine and metyrapone, indicating that the effect of atrazine was consistent across life stages and levels of the metyrapone treatment. Thus, these effects were dropped from the statistical model. The resulting simplified model revealed that frogs with early-life exposure to atrazine were less tolerant of *Bd* infections later in life than frogs that were not exposed to atrazine (atrazine×*Bd* load: Fi,34 = 4.50, *p* = 0.04; Fig. 5b). Corticosterone on experimental day 6 was not correlated significantly with resistance or tolerance of infections *(p* > 0.05, see Supplemental Table 1).

**Fig. 5.**
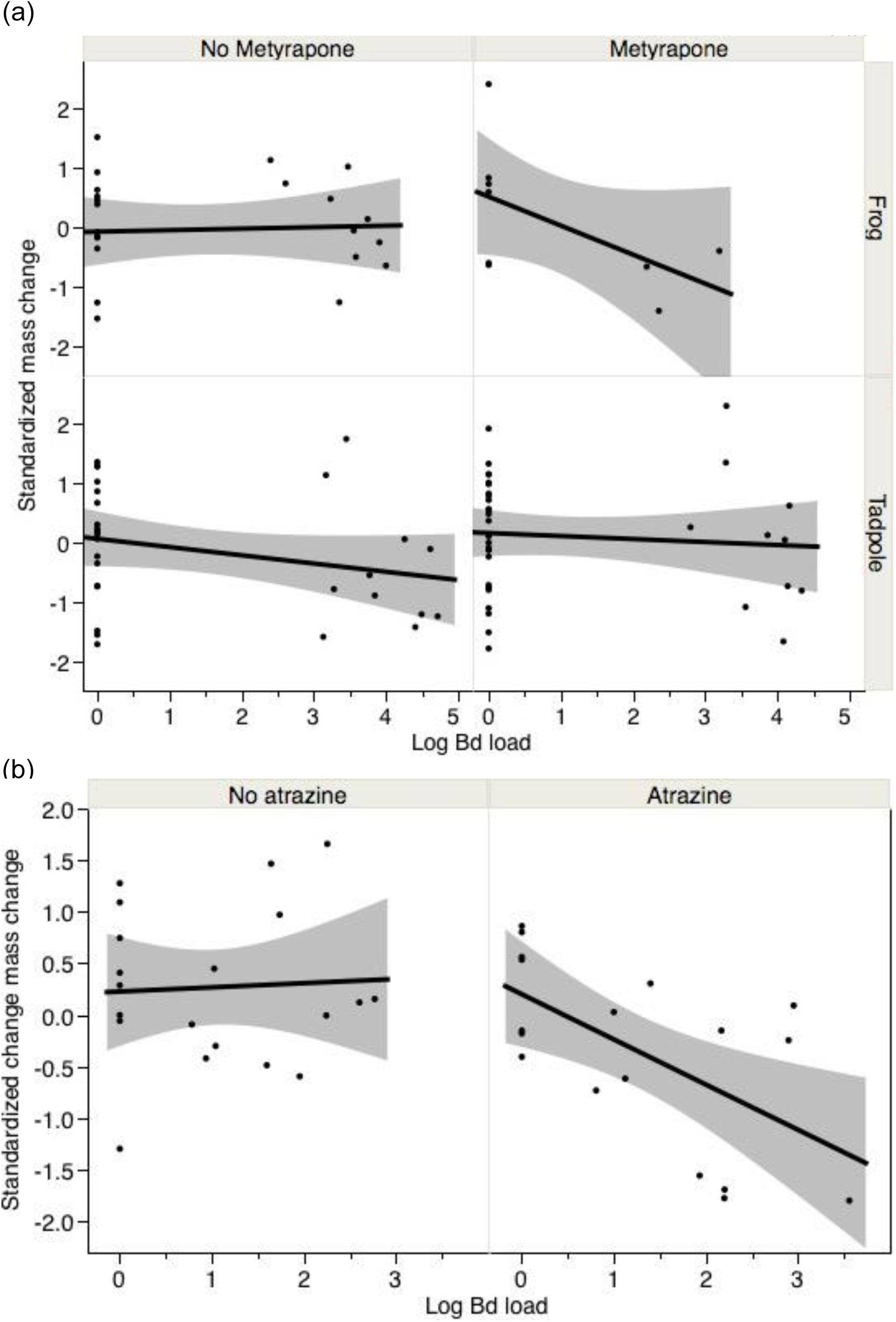
The relationship between tolerance (standardized mass change) and log *Batrachochytrium dendrobatidis (Bd)* load for *Osteopilus septentrionalis:* (a) 15 days (day 21) after *Bd* exposure pre-metamorphosis and five weeks after *Bd* exposure post-metamorphosis (all individuals were previously exposed to metyrapone (corticosterone synthesis inhibitor) for six days), and (b) six days after atrazine exposure in tadpoles. The life stage×metyrapone×*Bd* load interaction in panel (a) and the atrazine×*Bd* load interactions in panel (b) are significant *(p* < 0.05). See Results for statistics.

### Discussion

We explored whether the negative effects of atrazine and *Bd* on growth, development, and survival of Cuban treefrogs were mediated by the stress hormone corticosterone. We found that corticosterone levels in Cuban treefrogs were elevated after exposure to both an ecologically relevant concentration of atrazine and the fungal pathogen *Bd.* Importantly, exposure to metyrapone prevented these elevations in corticosterone (Fig. 2), demonstrating that it was successful in inhibiting corticosterone synthesis. However, we found little support for the hypothesis that the adverse effects of atrazine and *Bd* on growth, development, and tolerance to infection were mediated by corticosterone because corticosterone was not significantly correlated with any of these response variables (Supplemental Table 1) and because the observed effects of atrazine and *Bd* were not counteracted by exposure to metyrapone. Further studies are required to determine whether atrazine, *Bd*, and their effects on corticosterone mediate other events that we did not measure, such as metabolic regulation and oxidative balance.

Similar to our findings, several previous studies have shown that atrazine exposure can disrupt the HPA axis of vertebrates, dysregulating the production of the vertebrate stress hormones cortisol and corticosterone. For example, several studies showed that exposure to ecologically relevant concentrations of atrazine was associated with an increase in circulating corticosterone of small mammals (Fraites et al. 2009; Pruett et al. 2009; Riffle et al. 2014; Rogers et al. 2014) and cortisol of fish (Cericato et al. 2009; Koakoski et al. 2014). Exposure of salamanders and frogs to atrazine increased their circulating corticosterone levels (Larson et al. 1998; McMahon et al. 2017). Hernandez et al. (2014) showed that atrazine competitively inhibits corticosterone from binding with corticosterone binding globulin in both amphibians and mammals, further indicating that atrazine disrupts corticosterone regulation in these two vertebrate groups.

Similar to atrazine, *Bd* elevated corticosterone in our study, and this result is consistent with several previous studies demonstrating that infections can elevate stress hormones such as corticosterone in amphibians (Gabor et al. 2015; Peterson et al. 2013; Searle et al. 2014). Elevated corticosterone levels have also been found in wood frog tadpoles *(Rana sylvatica*) infected with ranavirus (Warne et al. 2011), and in lizards infected with parasites compared to non-infected controls (Dunlap and Schall 1995; Oppliger et al. 1998). However, some studies failed to find an effect of infections on circulating corticosterone in birds and frogs (Eggert et al. 2010; Kindermann et al. 2012; Knutie et al. 2013). Importantly, none of these studies experimentally tested whether corticosterone was mediating these responses to disease by inhibiting corticosterone synthesis.

The corticosterone synthesis inhibitor that we used, metyrapone, had different effects than atrazine on survival and development. Similar to Rohr et al. (2013), we did not find an effect of atrazine on short-term or long-term survival of Cuban treefrogs. Additionally, we found that early-life exposure to atrazine reduced growth rates, consistent with several previous studies (Rohr and McCoy 2010; Rohr and Palmer 2013). However, this effect on body size before metamorphosis was not significant after metamorphosis. In contrast, metyrapone had no significant effect on survival or body size before metamorphosis, close to when the actual metyrapone exposures occurred, but significantly decreased survival and body size after metamorphosis (Fig. 3b,c). Thus, the effects of metyrapone were persistent but delayed to later in life. These effects are likely caused by either the direct effect of metyrapone exposure or the indirect effect of low baseline levels of corticosterone on growth and survival.

We also note that metyrapone did not reduce corticosterone in the absence of stressors. This might be because the non-stressed tadpoles (such as the controls) could already be approaching the lower bound levels of baseline corticosterone necessary for homeostasis or because metyrapone cannot reduce circulating levels of corticosterone, only new corticosterone synthesis. Thus, in the absence of a stressor, we might not expect metyrapone to reduce baseline levels of circulating corticosterone if the half-life of corticosterone is reasonably long (see Supplemental material, Discussion of corticosterone half-lives). Similar to our finding, Glennemeier and Denver (2002b) found that metyrapone lowered corticosterone when individuals were stressed but not when they were not stressed.

We found that both metyrapone and atrazine affected amphibian defenses against *Bd.* Metyrapone exposure was associated with reduced tolerance to *Bd* when infections occurred after, but not before, metamorphosis (Fig. 5a), which is consistent with the delayed effects of metyrapone on growth and survival. While it is possible that reduced tolerance to *Bd* was mediated by dysregulation of corticosterone during *Bd* exposure, this finding is more likely caused by the accumulation of negative effects of metyrapone exposure through time and differences in susceptibility to *Bd* between the two life stages (Fisher et al. 2009; Rohr et al. 2013). Unlike our finding for metyrapone, we found that atrazine lowered tolerance of *Bd* infections similarly for both tadpoles and post-metamorphic frogs (Fig. 5b), results that match those of Rohr et al. (2013). Early-life exposure to atrazine reduced mass in Bd-infected tadpoles and post-metamorphic frogs, indicating that early-life exposure to atrazine had both short-and long-term effects on tolerance to *Bd*.

Although atrazine reduced mass in *Bd*-infected individuals of both life stages, exposure to metyrapone did not significantly counter these reductions in *Bd* tolerance, and thus, atrazine-induced elevations in corticosterone levels could not account for atrazine-induced reductions in tolerance of *Bd*. Although metyrapone had adverse effects later in life, these adverse effects could not account for the fact that metyrapone did not counteract the adverse effects of atrazine early in life when it did not have detectable effects and thus it seems unlikely that any adverse effects of metyrapone could be masking any corticosterone-mediated effects. Given that corticosterone does not appear to primarily mediate the effects of atrazine or *Bd* on amphibian growth or development, or the effects of atrazine on tolerance of *Bd* infections, these effects are more likely caused by direct effects of atrazine and *Bd* exposure, such as through energy lost from atrazine detoxification, defense against *Bd*, or repair from damage caused by atrazine or *Bd* (McMahon et al. 2013b; Voyles et al. 2009). Alternatively, these effects could be from indirect effects of atrazine and *Bd* on unmeasured hormones, such as thyroxine or steroidal sex hormones. For example, in larval tiger salamanders, atrazine exposure elevated thyroxine, another hormone associated with amphibian growth and metamorphosis (Larson et al. 1998). Additionally, several meta-analyses have revealed a negative association between testosterone and immunity, with sexually mature male vertebrates often exhibiting greater susceptibility to infection and higher parasite burdens in the field (Zuk 1996; Zuk and McKean 1996). Moreover, recent studies have revealed a causal relationship between testosterone, reduced immunity, and increased parasite loads in mice (Krucken et al. 2005; Lotter et al. 2013).

In contrast to the effects of atrazine on *Bd* tolerance, the effect of atrazine on resistance to *Bd* infections (i.e., lowered *Bd* load) did depend significantly on exposure to metyrapone (Fig. 4b). Resistance to *Bd* was higher when frogs were exposed to atrazine alone than to a combination of atrazine and metyrapone. However, our data do not strongly support the hypothesis that corticosterone was mediating this altered resistance to *Bd*. First, the corticosterone patterns associated with the atrazine treatment in the presence and absence of metyrapone (Fig. 2a) are not parallel to the patterns of resistance across atrazine and metyrapone treatments (Fig. 4b). Second, we did not find a significant interaction between atrazine, metyrapone, and *Bd* exposure on corticosterone, suggesting that these together did not mediate the observed resistance pattern. Third, there was no significant correlation between day 6 corticosterone and *Bd* abundance on tadpoles and post-metamorphic frogs (Supplemental Table 1). There was also no significant correlation between day 13, 14 corticosterone and *Bd* abundance on tadpoles (Pearson’s R = - 0.070; *p* = 0.70) or post-metamorphic frogs (Pearson’s R = 0.040; *p* = 0.87). Hence, our results regarding the relationship between corticosterone and *Bd* are equivocal, much like the literature on this topic. For example, some research suggests that corticosterone increases resistance to *Bd* (Murone et al. 2016; Tatiersky et al. 2015), whereas other research suggests that it has no effects (Searle et al. 2014). Additional work is needed to more thoroughly grasp the generality of effects of corticosterone on resistance to infections.

In conclusion, we found that there are costs of exposure to atrazine and *Bd*, which supports the results of other studies (Rohr and McCoy 2010; Rohr et al. 2013). However, by inhibiting corticosterone production with metyrapone, we found that increased corticosterone from atrazine and *Bd* exposure may not be the main factor mediating the observed decreases in growth, development, and tolerance to infection in Cuban treefrogs. Instead, our findings might largely be caused by repair from any damage caused by atrazine and *Bd* or indirect effects of atrazine and *Bd* on hormones other than corticosterone. Increased exposure to contaminants and pathogens are just two of many examples of how human activities are adversely affecting biodiversity and more studies are required to understand the mechanisms driving these effects (Dirzo et al. 2014). In particular, given that measurements of stress hormones are regularly being used to direct the management of wildlife populations (reviewed by Busch and Hayward 2009; Sheriff et al. 2011), additional studies are required to evaluate whether the adverse effects of the Anthropocene on biodiversity are often mediated by the effects of anthropogenic factors on the stress responses of organisms and their subsequent impacts on fitness.

## Acknowledgements

We thank J. Middlemis Maher for help with using metyrapone, E. Sauer for help with preparing *Bd* for inoculations, and N. Halstead for help with atrazine methodology, and early discussions with L. Martin and R. Boughton on this topic. We thank R. Earley for helpful discussion on the experimental design. We also thank K. Cunningham for measuring tadpoles, J. Reyes for help running hormone plates, M. Ehrsam for help with feeding and recording behavior, A. Dubour and V. Caponera for help with water-borne hormone collection, D. Pike for help with swabbing tadpoles for *Bd*, and S. Sehgal and S. Peters for help with animal husbandry.

## Compliance with ethical standards

### Funding

C.R.G. was funded by a REP grant from Texas State University. J.R.R. was funded by the National Science Foundation (EF-1241889), National Institutes of Health (R01GM109499, R01TW010286), US Department of Agriculture (NRI 2006-01370, 2009-35102-0543), and US Environmental Protection Agency (CAREER 83518801).

### Ethical approval

All applicable institutional and/or national guidelines for the care and use of animals were followed. This project was approved by the Texas State University Animal Care and Use Committee # 201485314.

## References

Boekelheide K et al. (2012) Predicting later-life outcomes of early-life exposures. Environ Health Perspect 120:1353–1361. doi:10.1289/ehp.1204934

Boonstra R (2013) The ecology of stress: a marriage of disciplines. Funct Ecol 27:7–10. doi:10.1111/1365-2435.12048

Boyle D, Boyle D, Olsen V, Morgan J, Hyatt A (2004) Rapid quantitative detection of chytridiomycosis (*Batrachochytrium dendrobatidis*) in amphibian samples using real-time Taqman PCR assay. Dis Aquat Org 60:141

Busch DS, Hayward LS (2009) Stress in a conservation context: A discussion of glucocorticoid actions and how levels change with conservation-relevant variables. Biol Conserv 142:2844–2853. doi:10.1016/j.biocon.2009.08.013

Cericato L et al. (2009) Responsiveness of the interrenal tissue of Jundia (*Rhamdia quelen*) to an in vivo ACTH test following acute exposure to sublethal concentrations of agrichemicals. Comp Biochem Physiol C Toxicol Pharmacol 149:363–367. doi:10.1016/j.cbpc.2008.09.002

Denver RJ (2009) Stress hormones mediate environment-genotype interactions during amphibian development. Gen Comp Endocrinol 164:20–31. doi:10.1016/j.ygcen.2009.04.016

Dirzo R, Young HS, Galetti M, Ceballos G, Isaac NJ, Collen B (2014) Defaunation in the Anthropocene. Science 345:401–406. doi:10.1126/science.1251817

Dunlap KD, Schall JJ (1995) Hormonal alterations and reproductive inhibition in male fence lizards (*Sceloporus occidentalis*) infected with the malarial parasite *Plasmodium mexicanum*. Physiol Zool 68:608–621

Eggert LMF, Jodice PGR, O'Reilly KM (2010) Stress response of brown pelican nestlings to ectoparasite infestation. Gen Comp Endocrinol 166:33–38. doi:10.1016/j.ygcen.2009.08.009

Ellis T, James JD, Stewart C, Scott AP (2004) A non-invasive stress assay based upon measurement of free cortisol released into the water by rainbow trout. J Fish Biol 65:1233–1252. doi:10.1111/j.1095-8649.2004.00499.x

Fisher MC, Garner TWJ, Walker SF (2009) Global emergence of *Batrachochytrium dendrobatidis* and amphibian chytridiomycosis in space, time, and host. Annu Rev Microbiol 63:291–310. doi:doi:10.1146/annurev.micro.091208.073435

Fraites MJP, Cooper RL, Buckalew A, Jayaraman S, Mills L, Laws SC (2009) Characterization of the hypothalamic-pituitary-adrenal axis response to atrazine and metabolites in the female rat. Toxicol. Sci. 112:88–99. doi:10.1093/toxsci/kfp194

Gabor CR, Fisher MC, Bosch J (2015) Elevated corticosterone levels and changes in amphibian behavior are associated with *Batrachochytrium dendrobatidis (Bd*) infection and *Bd* lineage. PLoS ONE 10:e0122685. doi:10.1371/journal.pone.0122685

Gabor CR et al. (2016) A non-invasive water-borne assay of stress hormones in aquatic salamanders. Copeia 2016:172–181

Glennemeier KA, Denver RJ (2002a) Small changes in whole-body corticosterone content affect larval *Rana pipiens* fitness components. Gen Comp Endocrinol 127:16–25

Glennemeier KA, Denver RJ (2002b) Role for corticoids in mediating the response of *Rana pipiens* tadpoles to intraspecific competition. J Exp Zool 292:32–40. doi:10.1002/jez.1140

Glennemeier KA, Denver RJ (2002c) Developmental changes in interrenal responsiveness in anuran amphibians. Integr Comp Biol 42:565–573. doi:10.1093/icb/42.3.565

Gosner K (1960) A simplified table for staging anuran embryos and larvae with notes on identification. Herpetologica 16:183–190

Hayes T, Falso P, Gallipeau S, Stice M (2010) The cause of global amphibian declines: a developmental endocrinologist's perspective. J Exp Biol 213:921 –933

Hernandez SE, Sernia C, Bradley AJ (2014) Effect of atrazine and fenitrothion at no-observed-effect-levels (NOEL) on amphibian and mammalian corticosterone-binding-globulin (CBG). Toxicol. Lett. 230:408–412. doi:10.1016/j.toxlet.2014.08.015

Hossie TJ, Ferland-Raymond B, Burness G, Murray DL (2010) Morphological and behavioural responses of frog tadpoles to perceived predation risk: A possible role for corticosterone mediation? Ecoscience 17:100–108. doi:10.2980/17-1-3312

Jones DK et al. (2017) Effect of simultaneous amphibian exposure to pesticides and an emerging fungal pathogen, *Batrachochytrium dendrobatidis*. Environ Sci Technol 51:671–679. doi:10.1021/acs.est.6b06055

Kiely T, Donaldson D, Grube A (2004) Pesticide industry sales and usage: 2000 and 2001 market estimates. U.S. Environmental Protection Agency, Washington, D.C.

Kindermann C, Narayan EJ, Hero J-M (2012) Urinary corticosterone metabolites and chytridiomycosis disease prevalence in a free-living population of male Stony Creek frogs (*Litoria wilcoxii*). Comp Biochem Physiol A 162:171–176

Knutie SA, Koop JAH, French SS, Clayton DH (2013) Experimental test of the effect of introduced hematophagous flies on corticosterone levels of breeding Darwin's finches. Gen Comp Endocrinol 193:68–71. doi:10.1016/j.ygcen.2013.07.009

Koakoski G et al. (2014) Agrichemicals chronically inhibit the cortisol response to stress in fish. Chemosphere 112:85–91. doi:10.1016/j.chemosphere.2014.02.083

Krucken J et al. (2005) Testosterone suppresses protective responses of the liver to blood-stage malaria. Infect Immun 73:436–443. doi:10.1128/iai.73.1.436-443.2005

Larson DL, McDonald S, Fivizzani AJ, Newton WE, Hamilton SJ (1998) Effects of the herbicide atrazine on *Ambystoma tigrinum* metamorphosis: duration, larval growth, and hormonal response. Physiol Zool 71:671–679

Lotter H et al. (2013) Testosterone increases susceptibility to amebic liver abscess in mice and mediates inhibition of IFN? Secretion in natural killer T cells. PLoS ONE 8:e55694. doi:10.1371/journal.pone.0055694

Martin LB, Hopkins WA, Mydlarz LD, Rohr JR (2010) The effects of anthropogenic global changes on immune functions and disease resistance. In: Ostfeld RS, Schlesinger WH (eds) Year in Ecology and Conservation Biology, vol 1195, pp 129–148

Matthews SG (2002) Early programming of the hypothalamo-pituitary-adrenal axis. Trends Endocrinol Metab 13:373–380

McMahon TA, Romansic JM, Rohr JR (2013a) Nonmonotonic and monotonic effects of pesticides on the pathogenic fungus *Batrachochytrium dendrobatidis* in culture and on tadpoles. Environ Sci Technol 47:7958–7964. doi:10.1021/es401725s

McMahon TA et al. (2013b) Chytrid fungus *Batrachochytrium dendrobatidis* has nonamphibian hosts and releases chemicals that cause pathology in the absence of infection. P. Natl. Acad. Sci. USA 110:210–215. doi:10.1073/pnas.1200592110

McMahon TA et al. (2014) Amphibians acquire resistance to live and dead fungus overcoming fungal immunosuppression. Nature 511:224–227. doi:10.1038/nature13491

McMahon TA, Boughton RK, Martin LB, Rohr JR (2017) Exposure to the Herbicide Atrazine Nonlinearly Affects Tadpole Corticosterone Levels. J Herpetol 51:270–273. doi:10.1670/16-126

Middlemis Maher J, Werner EE, Denver RJ (2013) Stress hormones mediate predator-induced phenotypic plasticity in amphibian tadpoles. Proc R Soc Biol Sci Ser B 280:20123075. doi:10.1098/rspb.2012.3075

Murone J, DeMarchi JA, Venesky MD (2016) Exposure to corticosterone affects hostresistance, but not tolerance, to an emerging fungal pathogen. PLoS ONE 11:e0163736. doi:10.1371/journal.pone.0163736

Neuman-Lee LA, Stokes AN, Greenfield S, Hopkins GR, Brodie ED, Jr., French SS (2015) The role of corticosterone and toxicity in the antipredator behavior of the Rough-skinned Newt (*Taricha granulosa*). Gen Comp Endocrinol 213:59–64. doi:10.1016/j.ygcen.2014.12.006

Oppliger, Clobert, Lecomte, Lorenzon, Boudjemadi, John A (1998) Environmental stress increases the prevalence and intensity of blood parasite infection in the common lizard *Lacerta vivipara*. Ecol Lett 1:129–138. doi:10.1046/j.1461-0248.1998.00028.x

Peterson JD et al. (2013) Host stress response is important for the pathogenesis of the deadly amphibian disease, chytridiomycosis, in *Litoria caerulea*. PLoS ONE 8. doi:e62146 1371/journal.pone.0062146

Pruett SB, Fan R, Zheng Q, Schwab C (2009) Patterns of immunotoxicity associated with chronic as compared with acute exposure to chemical or physical stressors and their relevance with regard to the role of stress and with regard to immunotoxicity testing. Toxicol. Sci. 109:265–275. doi:10.1093/toxsci/kfp073

R Core Team (2013) R: A language and environment for statistical computing. R Foundation for Statistical Computing, Vienna, Austria)

Råberg L, Sim D, Read AF (2007) Disentangling genetic variation for resistance and tolerance to infectious diseases in animals. Science 318:812

Råberg L, Graham AL, Read AF (2009) Decomposing health: tolerance and resistance to parasites in animals. Philos Trans R Soc B 364:37–49. doi:10.1098/rstb.2008.0184

Riffle BW et al. (2014) Novel molecular events associated with altered steroidogenesis induced by exposure to atrazine in the intact and castrate male rat. Reprod. Toxicol. 47:59–69. doi:10.1016/j.reprotox.2014.05.008

Rogers JM et al. (2014) Elevated blood pressure in offspring of rats exposed to diverse chemicals during pregnancy. Toxicol. Sci. 137:436–446. doi:10.1093/toxsci/kft248

Rohr JR et al. (2004) Multiple stressors and salamanders: Effects of an herbicide, food limitation, and hydroperiod. Ecol Appl 14:1028–1040. doi:10.2307/4493602

Rohr JR, Raffel TR, Hall CA (2010) Developmental variation in resistance and tolerance in a multi-host–parasite system. Funct Ecol 24:1110–1121. doi:10.1111/j.1365-2435.2010.01709.x

Rohr JR, McCoy KA (2010) A qualitative meta-analysis reveals consistent effects of atrazine on freshwater fish and amphibians. Environ Health Perspect 118:20–32. doi:10.1289/ehp.0901164

Rohr JR et al. (2013) Early-life exposure to a herbicide has enduring effects on pathogen-induced mortality. Proc R Soc Biol Sci Ser B 280:20131502–20131502. doi:10.1098/rspb.2013.1502

Rohr JR, Palmer BD (2013) Climate change, multiple stressors, and the decline of ectotherms. Conserv Biol 27:741–751. doi:10.1111/cobi.12086

Rollins-Smith LA (1998) Metamorphosis and the amphibian immune system. Immunol Rev 166:221–230. doi:10.1111/j.1600-065X.1998.tb01265.x

Rollins-Smith LA, Ramsey JP, Pask JD, Reinert LK, Woodhams DC (2011) Amphibian immune defenses against chytridiomycosis: impacts of changing environments. Integr Comp Biol 51:552–562. doi:10.1093/icb/icr095

Searle CL, Belden LK, Du P, Blaustein AR (2014) Stress and chytridiomycosis: exogenous exposure to corticosterone does not alter amphibian susceptibility to a fungal pathogen. J Exp Zool A Ecol Genet Physiol 321. doi:10.1002/jez.1855

Sheriff MJ, Dantzer B, Delehanty B, Palme R, Boonstra R (2011) Measuring stress in wildlife: techniques for quantifying glucocorticoids. Oecologia 166:869–887. doi:10.1007/s00442-011-1943-y

Stuart SN et al. (2004) Status and trends of amphibian declines and extinctions worldwide. Science 306:1783–1786

Tatiersky L et al. (2015) Effect of glucocorticoids on expression of cutaneous antimicrobial peptides in northern leopard frogs (*Lithobates pipiens*). BMC Veterinary Research 11:1–8. doi:10.1186/s12917-015-0506-6

Voyles J et al. (2009) Pathogenesis of chytridiomycosis, a cause of catastrophic amphibian declines. Science 326:582–585. doi:10.1126/science.1176765

Wake DB, Vredenburg VT (2008) Are we in the midst of the sixth mass extinction? A view from the world of amphibians. Proc Natl Acad Sci USA 105:11466–11473. doi:10.1073/pnas.0801921105

Warne RW, Crespi EJ, Brunner JL (2011) Escape from the pond: stress and developmental responses to ranavirus infection in wood frog tadpoles. Funct Ecol 25:139–146. doi:10.1111/j.1365-2435.2010.01793.x

Zuk M (1996) Disease, endocrine-immune interactions, and sexual selection. Ecology 77:1037–1042. doi:10.2307/2265574

Zuk M, McKean KA (1996) Sex differences in parasite infections: Patterns and processes. Int. J. Parasitol. 26:1009–1023. doi:10.1016/s0020-7519(96)80001-4

